# Patterns of information segregation during working memory and attention revealed by dual-task interference in behavior, pupillometry, and EEG

**DOI:** 10.1101/2021.04.20.440675

**Authors:** Justin T. Fleming, J. Michelle Njoroge, Abigail L. Noyce, Tyler K. Perrahione, Barbara G. Shinn-Cunningham

**Author notes:** Corresponding author: Justin T. Fleming. The authors declare no competing interests.

## Abstract

Making sense of our environment requires us to extract simultaneous temporal and spatial information from multiple sensory modalities, particularly audition and vision. This sensory information can be stored in working memory (WM) to guide future actions, at which point it must be safeguarded against interference from ongoing sensory processing. Recent fMRI research has uncovered regions in human frontal cortex well-suited to coordinate this interplay between attention and WM for multisensory and multidimensional information. Which of these brain regions are engaged depends on both the sensory modality of the input and the information domain of the task, forming the basis of two complementary networks specialized for auditory/temporal and visual/spatial processing. Motivated by the functional specializations of these networks, we examined whether similarity in sensory modality and information domain modulates neural and perceptual interference between two concurrent tasks. Participants stored temporal or spatial information about auditory or visual stimuli in WM, and on some trials, performed an intervening temporal or spatial auditory task during WM retention. WM recall and auditory perceptual judgments were impaired when the two tasks relied on the same sensory modality and/or information domain. Pupil dilations were also larger in these conditions, indicating increased cognitive effort. Event-related potentials (ERPs) revealed a neural signature of domain-based interference that was masked by behavioral ceiling effects. These results demonstrate that modality and information domain jointly affect how task information is represented in WM, and concomitantly, how tasks engage the complementary auditory-temporal and visual/spatial cognitive control networks.

## Introduction

Navigating everyday environments requires us to process spatial and temporal information from multiple sensory modalities, particularly vision and audition. Once stored in working memory (WM), these perceptual inputs can guide future actions. For instance, when crossing a busy street, we first look in one direction, sampling the spatial positions of vehicles over time to determine their direction and speed of movement. This information then moves into WM while we look the other way to determine the paths of vehicles approaching from that direction. To determine when it is safe to cross, we must maintain the temporal and spatial information stored in WM while simultaneously processing additional sensory inputs.

Under ideal circumstances, ongoing perceptual processing would not disrupt information stored in WM, and vice versa. Unfortunately, interference between WM maintenance and a concurrent task occurs all the time, especially when the information in WM and the sensory inputs from a secondary task share common attributes. For example, participants experience increased word-color Stroop interference when WM is loaded with similar verbal information, but not when WM is loaded with unrelated spatial information (Kim et al., 2005). Other studies featuring a WM task and distractor stimuli have shown that distractors selectively interfere with recall of closely related items from WM. For instance, in visual-spatial WM, distractors that vary in polar angle selectively impair recall of a target polar angle (Marini et al., 2017); in verbal WM, distractor words selectively interfere with participants’ ability to recall phonologically similar target words (Oberauer & Lange, 2008). These findings support the “multiple resource theory” account of dual-task interference, which posits that two tasks are more likely to interfere with each other when they draw on a shared pool of specific cognitive resources (Navon & Gopher, 1979; Nickerson, 1980; Wickens, 2002).

Concurrent tasks can also interfere with one another if they require processing information from the same sensory modality. In dual-task paradigms, increased interference has been observed when both tasks are auditory, visual, or tactile compared to conditions in which the tasks operate on inputs from different sensory modalities (Morrison et al., 2015; Scerra & Brill, 2012). Additionally, auditory and visual discrimination thresholds (for pitch and contrast, respectively) are unaffected by concurrent, irrelevant stimuli in the opposite modality, but are considerably worse in the presence of concurrent stimuli in the same modality (Alais et al., 2006).

Recent fMRI research has identified subregions within human frontal cortex (LFC) that are preferentially driven by auditory or visual information during attention and WM tasks (Braga et al., 2017; Mayer et al., 2016; Michalka et al., 2015; Noyce et al., 2017). These LFC subregions are strongly connected to the corresponding primary sensory brain areas, forming networks tuned to processing either auditory or visual information and storing it in WM (Michalka et al., 2015; Tobyne et al., 2017). The organization of these networks may explain the patterns of interference based on sensory modality reported in previous studies; an auditory and a visual task should drive separate sensory-biased networks and therefore interfere relatively little with one another, whereas tasks presented in the same sensory modality should interfere more.

Although each LFC network is biased toward information from one sensory modality, each network can also be recruited to process information from the non-preferred sensory modality. This occurs in a manner consistent with the complementary specializations of vision for spatial processing and audition for temporal processing. From the retina, visual representations are inherently spatial, and neural maps of space are found throughout the visual processing pathway (Silver & Kastner, 2009; Stensaas et al., 1974; Swisher et al., 2007). In contrast, auditory spatial information must be indirectly computed by comparing the signals reaching the two ears. The peripheral auditory system instead excels at representing temporal information; each auditory nerve fiber, tuned to a narrow acoustic frequency range, phase-locks to the temporal structure of the portion of the sound pressure wave to which it responds (Dynes & Delgutte, 1992). Reflecting these complementary strengths, auditory tasks that involve *spatial* information processing recruit the visual-biased LFC network, while visual tasks that involve *temporal* judgments recruit the auditory-biased LFC network (Michalka et al., 2015). This flexible allocation of resources allows information from either sensory modality to be processed by a network specialized for the temporal and spatial strengths of audition and vision, respectively.

The functional specializations of these networks point to a potential neural substrate for the “resources” in multiple resource theory. In a dual-task setting (such as performing a perceptual task under WM load), two tasks that rely on the same LFC network should interfere strongly with each other, whereas tasks that rely on different networks should interfere less. In addition to explaining the greater interference between tasks in the same sensory modality, this pattern of cross-modal LFC network recruitment yields a previously untested prediction: that dual-task interference may depend on interactions between sensory modality *and* information domain (e.g., whether the task is temporal vs. spatial). For example, performing an auditory-temporal perceptual task should strongly interfere with auditory-temporal information stored in WM. On the other hand, the same perceptual task might interfere less with auditory-*spatial* WM; even though both tasks operate on auditory inputs, auditory-spatial information may be stored and maintained in the visual-biased network, resulting in less interference with auditory-temporal perceptual processing. The present work tests the idea that, in addition to similarity in sensory modality, interference between WM and perceptual tasks will be modulated by similarity in information domain. We developed a dual-task interference paradigm featuring WM and “Intervening” tasks. The WM task required participants to remember either temporal or spatial information about a set of nonverbal auditory or visual stimuli. The Intervening task, which was presented during WM retention, required participants to make an immediate perceptual judgment about either the timing or spatial locations of auditory stimuli. This paradigm allowed us to observe patterns of interference as participants engaged a WM network to encode sensory information in memory, performed a real-time perceptual task that required either increasing the load on the active network or switching networks, and finally retrieved the information stored in WM.

In addition to behavioral performance on the Intervening and WM tasks, we simultaneously measured pupillometry and electroencephalography (EEG) as participants encoded and held information in WM. Pupillometry serves as a sensitive psychophysiological index of the cognitive effort required to achieve a certain performance level, even in the absence of behavioral differences. For instance, maximum pupil dilations evoked by speech are larger for listeners with hearing loss than normal-hearing controls, even at high signal-to-noise ratios where both groups perform at ceiling (Ohlenforst et al., 2017). Under some circumstances, pupil dilations also correlate with participants’ subjective assessments of task difficulty (Koelewijn et al., 2015). As a time-series measure, pupillometry can also provide insights into *when* participants deployed effort over the course of each dual-task trial. The EEG data provided an even more temporally resolved signal, allowing us to examine the effects of dual-task interference on the neural responses to individual stimulus events (event-related potentials, ERPs) and an established oscillatory signature of WM maintenance (alpha power in the 8-13 Hz frequency band). Across these measures, we expected to see evidence of heightened interference when the Intervening and WM tasks were matched in sensory modality and/or information domain. We hypothesized that interference in these conditions would manifest as increased behavioral errors and larger task-evoked pupil dilations. Among the multiple possible effects that dual-task interference might have on EEG signals (Lin et al., 2011; Obleser et al., 2012; Serrien et al., 2004), we were particularly interested in testing the following predictions: As interference increased on the basis of sensory modality and information domain, ERP amplitudes would be reduced (reflecting taxation of resources shared between the two tasks) and ongoing alpha oscillations during WM maintenance would be interrupted.

## Materials and Methods

### Participants

Twenty-three healthy young adults completed all experimental procedures. Data from three participants were removed due to excessive noise in the pupillometry or EEG recordings, making for a final sample of *N*=20 (13 female; mean age 20.9 years; range 18 to 28 years). This recruitment target was based on a power analysis conducted on behavioral pilot data (N=6), which indicated that 15 participants would be required to detect differences in auditory perceptual performance based on the type of information held in WM. To account for variability between participants and the potential for additional noise in the EEG and pupillometry data, the recruitment goal was increased to 20.

One participant was excluded from only the time-frequency analyses due to anomalous high-frequency noise in their EEG data. All participants had normal or corrected-to-normal visual acuity and no reported colorblindness. Participants with corrected vision wore contact lenses instead of glasses to avoid potential artifacts in the pupillometry data. All participants had clinically normal hearing, defined by tone detection thresholds below 20 dB HL at octave frequencies between 250 Hz and 8 kHz, as confirmed by an audiometric screening. Participants gave written informed consent and were compensated for their participation. All study procedures were approved and overseen by the Boston University Charles River Campus Institutional Review Board.

### Experimental Setup

The experiment was conducted in a darkened, electrically shielded, sound-treated booth. Participants were seated comfortably with their chin resting on a desk-mounted head support (SR Research). A BenQ 1080p LED monitor (27-inch diagonal, 120 Hz refresh rate) was positioned in front of the participant at approximately 65 cm distance. The monitor was set to 3% of its maximum brightness level to prevent eye fatigue and pupil diameter saturation. An EyeLink 1000 Plus eye-tracking system was placed on the desk just below the display for measurement of pupil diameter. Six free-field loudspeakers (KEF E301) were mounted in an arc around the participant at a distance of 1.5 m. Five of the loudspeakers were equally spaced in azimuth at ±90°, ±45°, and 0° relative to midline; the sixth was placed immediately to the left of the central loudspeaker, at −4° azimuth, and used only in the Intervening tasks described below (Fig. 1A). All six loudspeakers were positioned at approximately 5° elevation relative to the horizontal plane of the eyes to reduce obstruction by the visual display. Auditory stimulus presentation was handled using an RME Fireface UCX soundcard.

**Figure 1:**
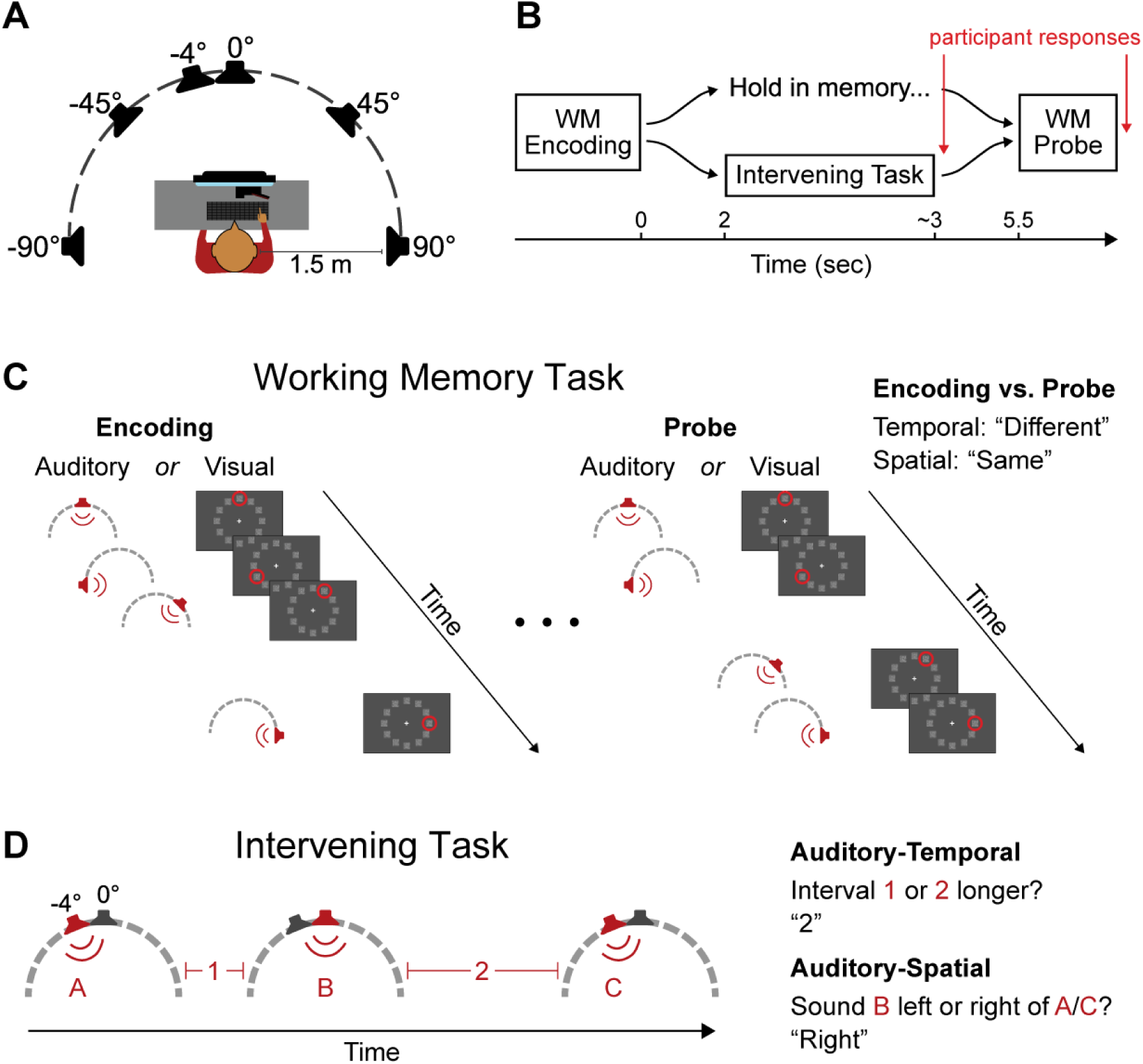
Experimental setup and task design. **A**, Overhead depiction of the experimental setup. All the loudspeakers except the one at −4° azimuth were used for the WM tasks, while only the loudspeakers at −4° and 0° were used for the Intervening tasks. **B**, Overall dual-task structure. The time base is relative to the offset of the final stimulus in the WM encoding phase. **C**, WM task structure. Loudspeakers playing each auditory stimulus and the changing visual stimuli are shown in red. Sensory modality always matched between the encoding and probe sequences. Correct responses for this example trial, shown on the right, differ depending on whether task was temporal or spatial. Note that in the actual experiment, inter-stimulus intervals were always isochronous in the auditory-spatial WM task. **D**, Intervening task structure. Auditory stimuli and inter-stimulus intervals are numbered and referred to in the temporal and spatial task responses on the right.

A standard keyboard was used to register all task responses. 64-channel EEG data were collected at a sampling rate of 2048 Hz using a Biosemi ActiveTwo system. Separate PCs were used for pupillometry recording and EEG recording, and a third PC was used for presenting stimuli and registering behavioral responses. To ensure synchrony of event triggers (e.g., trial starts, stimulus presentation) between the pupillometry and EEG data, triggers were output through the S/PDIF channel on the soundcard, converted to TTL pulses using a custom converter box, and written simultaneously into the EEG and pupillometry data files. Experiment control was carried out using custom MATLAB software, and visual stimulus presentation was implemented using the Psychtoolbox package (Brainard, 1997).

### Task and Experimental Design

Participants performed a dual-task paradigm, comprising a working memory (WM) task and an Intervening task (Fig. 1B). Each trial started with a 1.5-s baseline period, followed by the presentation of a sequence of four auditory or visual stimuli to be encoded in WM. Each stimulus was presented at one of five (auditory) or twelve (visual) locations, and each inter-stimulus interval in the sequence was randomly set to be either short or long (more details below). The WM task could be either temporal or spatial, yielding four total WM task conditions: auditory-temporal (AT), auditory-spatial (AS), visual-temporal (VT), and visual-spatial (VS). When the WM task was temporal, participants were instructed to remember the pattern of inter-stimulus intervals (i.e., the rhythm), regardless of spatial locations. When the WM task domain was spatial, participants had to remember the locations of the stimuli, regardless of order or timing (Fig. 1C). In all but the AS WM task (see below), both the locations and intervals could change between the encoding and probe sequences, but participants were instructed to ignore changes in the unattended domain. These WM task conditions were derived from tasks used to recruit the complementary LFC networks in fMRI studies (Michalka et al., 2015).

Participants retained stimulus information in WM for 5.5 seconds, after which a four-stimulus probe sequence was presented in the same sensory modality as the encoded sequence. Participants compared the encoded and probe sequences and made a same-different judgment on the remembered domain (temporal or spatial). After the conclusion of the probe stimulus, participants had 1.5 sec to indicate whether the encoded and probe sequences were the same (by pressing “1” on the keyboard) or different (by pressing “0”). Each block contained an equal number of same and different trials, ordered randomly. Participants maintained fixation on a small black cross (0.41° visual angle) at the center of the display throughout the trial.

We conducted pilot testing of the WM tasks to determine parameter settings that avoided ceiling or floor effects in behavioral measures. For participants to perceive the different inter-stimulus intervals equally well, a larger temporal separation was needed for the visual stimuli (200 and 580 ms) than the auditory stimuli (200 and 340 ms). Conversely, the visual-spatial task was too easy with only five stimulus locations, so the number of potential visual locations was increased to 12. Finally, participants struggled to perform the AS WM task when stimulus timing was variable; therefore, both encoding and probe stimuli in this condition were presented isochronously at the longer inter-stimulus interval.

On some trials, participants also performed an Intervening task during the WM retention period. This task was always auditory to allow pupil diameter to be measured in the absence of any visual stimulation, but like the WM tasks, it could be either temporal or spatial (AT or AS; see Fig. 1D). The stimulus structure was the same for the temporal and spatial variants. Starting 2 sec after the offset of the final stimulus in the WM encoding phase, a sequence of three auditory stimuli was presented. These stimuli were white noise bursts, acoustically distinct from the stimuli used in the auditory WM tasks (tone complexes) to prevent confusion between the tasks. One of the two intervals between the stimuli was randomly chosen to be slightly longer than the other. The precise intervals were jittered on each trial, with an average interval duration of 460 ms and a 90 ms average difference between the two intervals. The sounds were presented from the two near-frontal loudspeakers (−4° and 0° azimuth). The first stimulus played from one of these loudspeakers, chosen randomly and with equal probability; the second stimulus was played from the other loudspeaker, and the third was played from the same location as the first. In the temporal Intervening task, participants judged whether the first or second inter-stimulus interval was longer. In the spatial Intervening task, participants were asked to determine whether the second sound was to the left or right relative to the first and third sounds.

This stimulus design allowed physically identical auditory stimuli to be used for the spatial and temporal Intervening task conditions. However, it did introduce an asymmetry between conditions in the amount of information required to do the task. In the temporal Intervening task, participants needed to attend all three stimuli in order to compare the two inter-stimulus intervals, whereas in the spatial Intervening task, participants were often able to make their judgment on the second auditory stimulus by comparing its location to the first. This could result in a longer period of increasing pupil size in the temporal task, leading to larger peak pupil diameter. In addition, behavioral accuracy differed between the two Intervening tasks. Both of these differences could confound comparisons across the different Intervening tasks; however, our analyses focused mainly on comparisons across the four WM conditions *within* each Intervening task, which minimized the impact that potential differences in difficulty and behavioral strategy could have had on our main conclusions.

Participants registered Intervening task responses with a keypress immediately after the last Intervening task stimulus. Thus, neural signatures of motor planning and execution may be present in the EEG data near the end of the WM retention phase. However, we deemed this less detrimental than asking participants to hold their Intervening task responses until the end of the trial, as such a design would increase the WM load in trials with an Intervening task and complicate comparison with the no Intervening task conditions. Furthermore, any motor components in the EEG data should be present whenever there was an Intervening task, allowing for fair comparison between the different combinations of WM and Intervening tasks.

Trials were grouped into blocks of 20. Within each block, the WM and (when present) Intervening task conditions were held constant. At the start of each block, an instruction screen indicated the sensory modality and relevant domain (temporal or spatial) for both tasks in the upcoming block. Participants were allowed to take untimed breaks between blocks. Participants performed one block of each WM and Intervening task combination to be completed in that session before any conditions were repeated, and the same condition was not allowed to repeat in adjacent blocks. In total, participants performed 40 trials of each combination of WM task modality, WM task domain, and Intervening task condition.

Each complete dataset required three separate visits to the lab. The first session was reserved for consent, audiometric screening, and task practice. Participants practiced each variant of the WM and Intervening tasks in isolation until they understood the procedure, then performed three to five example trials of the full dual-task paradigm. Data collection for the actual experiment occurred in the two subsequent sessions, which were split based on the sensory modality of the WM tasks; auditory and visual WM tasks were performed on separate days. The order of these sessions was randomized and counterbalanced across participants.

### Stimulus Details

For the visual WM tasks, 12 stimuli were arranged in a circle centered on the fixation cross and shown on a constant dark grey background (2.51 cd/m^2^). Each stimulus was a square patch of visual noise, subtending 2.86° of visual angle and composed of a 30 x 30 grid of smaller squares. Each of these smaller squares was filled with a greyscale color between black and white, such that the average luminance across the patch was 5.85 cd/m^2^. The angular spacing between each patch was 30°, and the entire stimulus circle subtended 21.59° of visual angle. To equate display luminance and structure across tasks, these visual stimuli remained present but static throughout the auditory WM and Intervening tasks. To employ these stimuli in the visual WM tasks, the luminance of each small square in a given patch could be resampled; this made the visual patch appear to jitter without changing the average luminance across the patch.

For the auditory WM tasks, each stimulus was a 50-ms tonal chord consisting of 3 harmonically unrelated complex tones (fundamental frequencies of 422, 563, and 670 Hz); each complex was made up of its first nine harmonics with equal amplitude. The same tone complex was used for all auditory stimuli in the WM tasks. For the Intervening tasks, which were always auditory, the stimuli were one of five pre-generated, 50-ms bursts of noise bandpass filtered between 100 and 10,000 Hz. Identical noise tokens were used for all three stimuli within each Intervening task sequence. Both types of stimuli were relatively broadband, thus ensuring they provided rich and robust spatial localization cues. All auditory stimuli were ramped on and off with a 5 ms cosine-squared ramp to avoid spectral splatter and transient onset and offset artifacts.

In the visual WM tasks, the first stimulus to change in the encoding and probe sequences was always the one at top-center (12 o’clock). Similarly, in all auditory WM sequences, the first sound was presented from the central loudspeaker. This was done to equate the number of items stored in WM across the spatial and temporal WM tasks; with four stimuli in each sequence, there were three intervals to remember for the temporal tasks, and so the first stimulus location was held constant such that only three locations needed to be remembered for the spatial tasks.

### Behavioral Data Analysis

The primary behavioral metrics in this study were error rates on the Intervening and WM tasks. Behavioral performance was statistically analyzed using logistic mixed effects regression models. For Intervening task performance, the model included fixed-effect terms for the WM task condition (AT, AS, VT, or VS) and the Intervening task type (AT or AS), as well as the interactions between these terms. Random-effects terms were included to capture participant-specific intercepts and slopes for both predictor variables. The model was structured as follows:

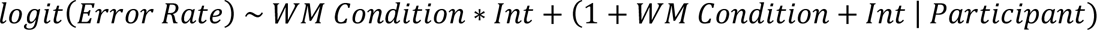

A similar model was used to analyze WM recall errors. To facilitate separately examining effects of WM task modality (auditory or visual) and domain (temporal or spatial), the WM condition was expressed as two fixed-effect terms in this model:

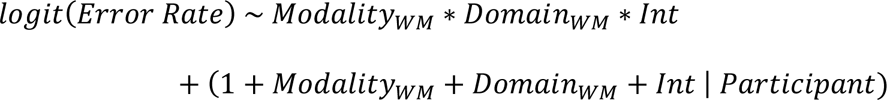

To investigate main effects and interactions at the group level, the coefficients from these models were fed into a two-way (Intervening task) or three-way (WM task) repeated measures ANOVA. P-values for model terms were estimated using the Satterthwaite approximation for degrees of freedom. Contrasts were treatment coded with baseline levels initially set as “auditory” for WM modality, “temporal” for WM domain, and “none” for the Intervening task. These choices had no impact on the outcome of the group-level ANOVA. Pairwise post-hoc testing was conducted by cycling which level was considered “baseline” for each factor until a β-weight (and corresponding p-value) could be extracted for each necessary pair of levels.

We did not attempt to precisely equate task difficulty across conditions for individual participants, which makes it difficult to compare performance across Intervening task or WM task conditions. Instead, we assessed how performance on one task (keeping the condition of that task fixed) changed across conditions of the other task. For Intervening task performance, this meant examining performance on each separate Intervening task as a function of the modality and domain of the WM task; for WM task performance, we examined recall in each separate WM condition as a function of the Intervening task condition. All post-hoc tests were corrected for multiple comparisons using the Bonferroni-Holm method, using the total number of comparisons made in each analysis. For example, the four WM task conditions led to six comparisons within each Intervening task, but the number of comparisons for correction was set to the total across both Intervening tasks (12).

### Pupillometry data collection and analysis

We measured task-evoked pupil dilations because they dynamically reflect the participant’s level of effort or arousal during task performance (Causse et al., 2016; Gilzenrat et al., 2012; Murphy et al., 2011; Winn et al., 2015). Prior to the start of each experimental session, eye position measurements were calibrated based on five fixation points (display center, ±20° azimuth on the horizontal plane, and ±10° elevation on the median plane). This calibration was validated at the center position prior to each trial, and at all five points at the start of each block. Whenever eye position was offset from the calibration by an average of greater than 4°, the full five-point calibration was repeated. The experimenter monitored gaze position throughout the experiment; if at any point during the trial the participant’s gaze deviated substantially from 0° in either azimuth or elevation, the experimenter immediately stepped in to ensure that the participant maintained center fixation moving forward.

A custom MATLAB analysis pipeline and wrapper GUI were used to prepare pupil data for statistical analysis. First, trials were split into baseline and trial windows, where the trial window spanned the start of the WM encoding sequence through the end of the WM retention phase. Next, blinks were automatically detected based on instantaneous position, velocity, and acceleration thresholds. An experimenter manually reviewed the data, and using the GUI, adjusted blink thresholds or manually marked additional blink segments as needed. Blinks and other marked segments of noisy data were replaced with a linear interpolation between the average of the three samples preceding and following the blink. When blinks occurred at the beginning of the trial window, a linear fit was made to the five samples following the blink, and this fit was back-projected through the blink segment. The opposite procedure was used for blinks falling at the end of the trial window. Trials in which more than 25% of the data was made up of rejected segments were automatically excluded from further analysis. When data from both eyes were available, the two traces were averaged. To produce the final output, the traces were concatenated, Z-scored, then split back into trials and trial windows. This procedure eschewed absolute pupil size measures in favor of values that were individually normalized for each participant, so only relative pupil diameter between conditions is interpreted.

Statistical testing for differences between conditions in the pupil time courses was carried out using non-parametric permutation tests. First, at each time point, the difference between two conditions was assessed parametrically using a paired T-test across the individual participant average data. Next, we created a null distribution for each participant by randomly shuffling the condition labels and performing the same analysis for 2000 iterations. These randomly labeled samples were used to recompute paired T-tests at each time point, generating a null distribution of 2000 such values. Finally, at each time point, the significance level of the observed difference was determined by calculating the percentage of the null distribution with a T-value equal to or larger than the T-value from the actual data. Significant differences were only considered reliable if the p-value fell below 0.05 for a minimum of 15 consecutive samples.

### EEG Data Analysis

EEG analyses were carried out using the FieldTrip package in MATLAB (Oostenveld et al., 2011). For event-related potential (ERP) analyses, EEG preprocessing comprised the following steps: read in the continuous data one channel at a time and immediately downsample to 256 Hz; re-reference the data to the average of two electrodes placed on the mastoids, bandpass filter between 0.5 and 20 Hz (zero-phase FIR filter, transition width of 0.2 Hz, order of 9274); manually identify and remove segments containing muscle artifacts; perform an independent components analysis (ICA) to project out blinks and saccadic eye movements; epoch the data from 100 ms before to 500 ms after each individual auditory or visual stimulus (timing differences between conditions precluded whole-trial averaging); reject any epochs in which the data exceeded a 100 µV peak-to-peak threshold; and baseline correct by subtracting off the mean of the first 100 ms of each epoch.

For time-frequency analyses, a similar preprocessing pipeline was used, but with a few key differences. First, the low-pass filter cutoff was raised to 80 Hz. Second, participant average ERPs (recomputed with the new filter cutoffs) were subtracted from the data in the time domain immediately prior to epoching; this step served to minimize phase-locked evoked contributions to the time-frequency response. Subtracted ERPs were specific to each trial phase (i.e. encoding, retention, and probe), WM and Intervening task condition, and position in the stimulus sequence. Third, the data were split into whole-trial epochs, spanning the baseline period through the final stimulus in the probe sequence, instead of shorter individual stimulus epochs. The continuous Morlet wavelet transform (wavelet width of 5 cycles in 1 Hz steps) was used to obtain the power spectra of each trial. Prior to wavelet analysis, the signal was padded to avoid edge artifacts. This was done by copying the first and last 5 seconds of the epoch, reflecting each copy on the time axis, then appending them to the beginning and end of the signal, respectively. Finally, the data were split into the three key trial phases: WM encoding, WM retention/Intervening task, and WM probe.

We extracted time courses from the resulting power spectra at each channel in the theta (4-8 Hz) and alpha (8-13 Hz) frequency bands. To produce theta time courses, power estimates centered on 4, 5, 6, and 7 Hz (1 Hz bandwith) were averaged for each participant. Individual differences in the peak frequency of alpha oscillations are well-established (Klimesch et al., 1999). Thus, for alpha time courses, the individual alpha frequency for each participant was defined as the frequency at which the absolute value of the alpha change relative to baseline was maximal during WM retention (the direction of alpha change was found to flip based on WM modality in the present study). Alpha power time courses were reported as the power at this frequency averaged with the power 1 Hz above and below the individually defined alpha frequency.

Similar to the pupillometry data, EEG data were analyzed using non-parametric permutation testing, this time implemented using FieldTrip software. To control the multiple comparisons problem with high-dimensional EEG data, we used cluster-based permutation testing to analyze ERPs and oscillatory power time courses (Maris & Oostenveld, 2007). Paired t-tests were first conducted at each time-channel data point. Those comparisons reaching a significance threshold (generally p < 0.01) were grouped into clusters of points contiguous in time and within 40 mm scalp distance of one another. The “mass” of this cluster was then computed as the sum of T-scores across all member points. Finally, this cluster mass was compared against a null distribution of clusters formed using the same analysis on 2000 random permutations of the condition labels.

## Results

### Intervening Task Performance

Performance accuracy on the Intervening tasks depended on the Intervening task condition (AT or AS) and its interaction with the type of information participants were holding in WM (AT, AS, VT, or VS; Fig. 2). These effects were analyzed using a logistic mixed-effects model, coefficients of which were then supplied to a Type-III ANOVA to determine the significance of effects at the group-level (see Materials and Methods). This ANOVA showed significant main effects of Intervening task (χ^2^(1,20) = 32.7, *p* < 0.0001) and WM task (χ^2^(3,20) = 17.6, *p* = 5.29·10^−4^), as well as a significant interaction between these factors (χ^2^(3,20) = 24.1, *p* < 0.0001). The main effect of Intervening task reflects overall lower error rates (better performance) on the AS compared to the AT Intervening task. The two loudspeakers used for the Intervening tasks were placed as close together as possible, but this spatial separation was nonetheless large enough for the AS Intervening task to be relatively easy for some participants. Given the overall performance difference between the two Intervening tasks, we examined each Intervening task separately in subsequent pairwise post-hoc tests; differences in these tests could only be caused by interference based on the type of information participants were holding in WM.

**Figure 2:**
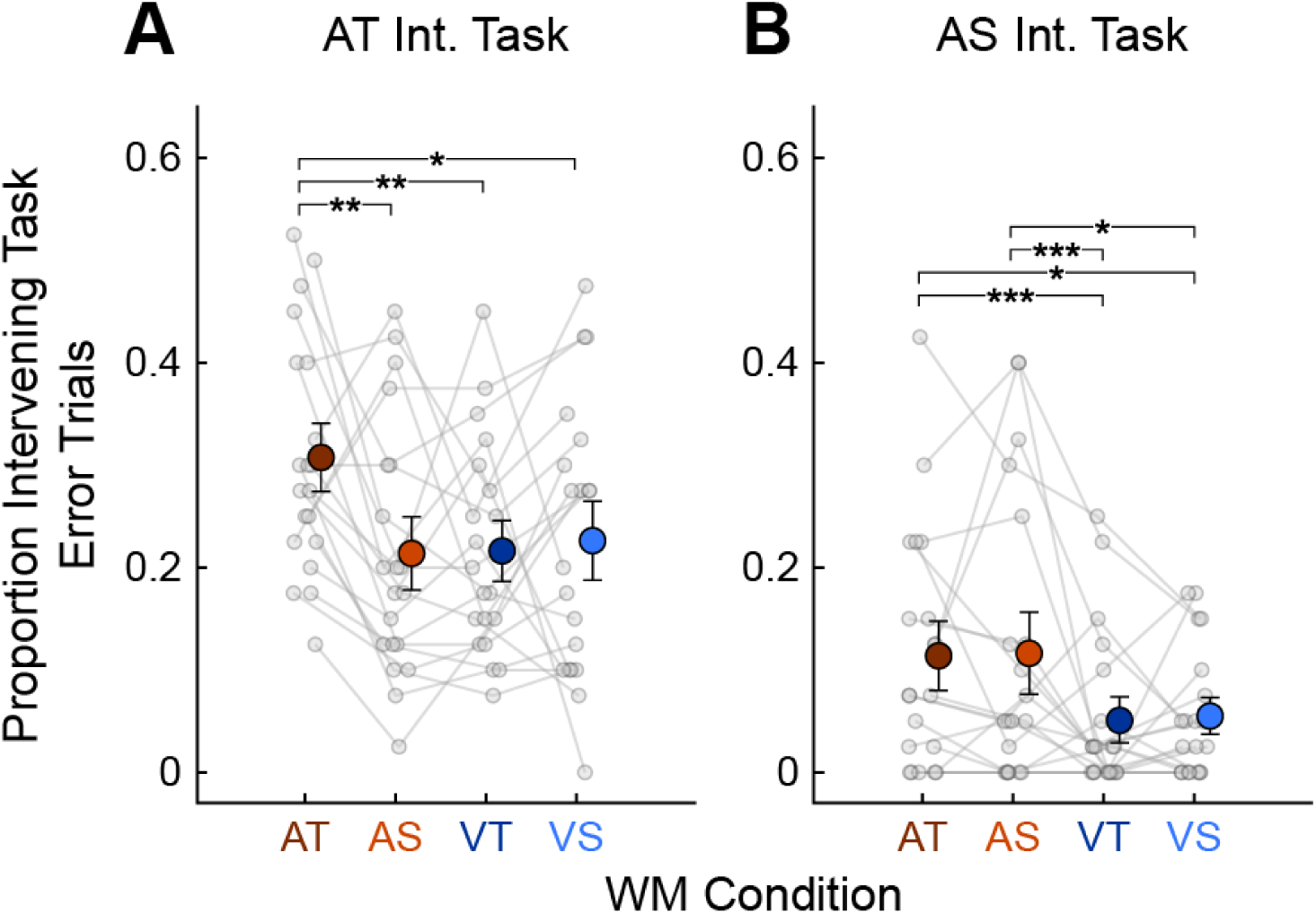
Intervening task error rates. Performance is shown for the AT (**A**) and AS (**B**) Intervening tasks. Grey points represent individual participants, large colored circles represent means, and error bars represent S.E.M. Chance performance is an error proportion of 0.5. Asterisks indicate significant differences between WM task levels in the mixed-effects model, examined separately within each Intervening task. * = *p* < 0.05, ** = *p* < 0.01, *** = *p* < 0.001 after Bonferroni-Holm correction.

Participants made significantly more errors on the AT Intervening task (Fig. 2A) when the information being held in WM was also AT, as compared to all the other WM conditions (AT vs. AS, *p* = 0.006; AT vs. VT, *p* = 0.003; AT vs. VS, *p* = 0.03). Accuracy on the AS Intervening task (Fig. 2B) appeared to be modulated primarily by the sensory modality of the WM task. Participants made significantly more errors when both tasks were auditory than when the WM task was visual (AT vs. VT, *p =* 1.76·10^−5^; AT vs. VS, *p =* 0.026; AS vs. VT, *p =* 5.50·10^−5^; AS vs. VS, *p =* 0.021). We also hypothesized that errors on the AS Intervening task might increase when the WM task was VS, as both tasks shared spatial processing demands, but this did not appear to be the case. However, the floor effect in AS Intervening task errors may have limited our ability to detect additional interference effects based on WM domain.

Importantly, these patterns of Intervening task performance cannot be explained by differences in task difficulty between the WM conditions. As will be shown below, participants made relatively few recall errors in the AT WM condition in the absence of an Intervening task, at least compared to the AS and VT WM conditions (see Fig. 3A below). If Intervening task performance varied mainly with the difficulty of the WM task, one would therefore expect relatively little interference in the AT WM condition, which in fact caused a relatively high degree of interference in both Intervening tasks. Instead, the patterns here reflect relationships between the types of processing demanded by the two tasks. Participants tended to make more errors on the Intervening task when its sensory modality and (in the case of the AT Intervening task) its information domain matched those of the information stored in WM.

**Figure 3:**
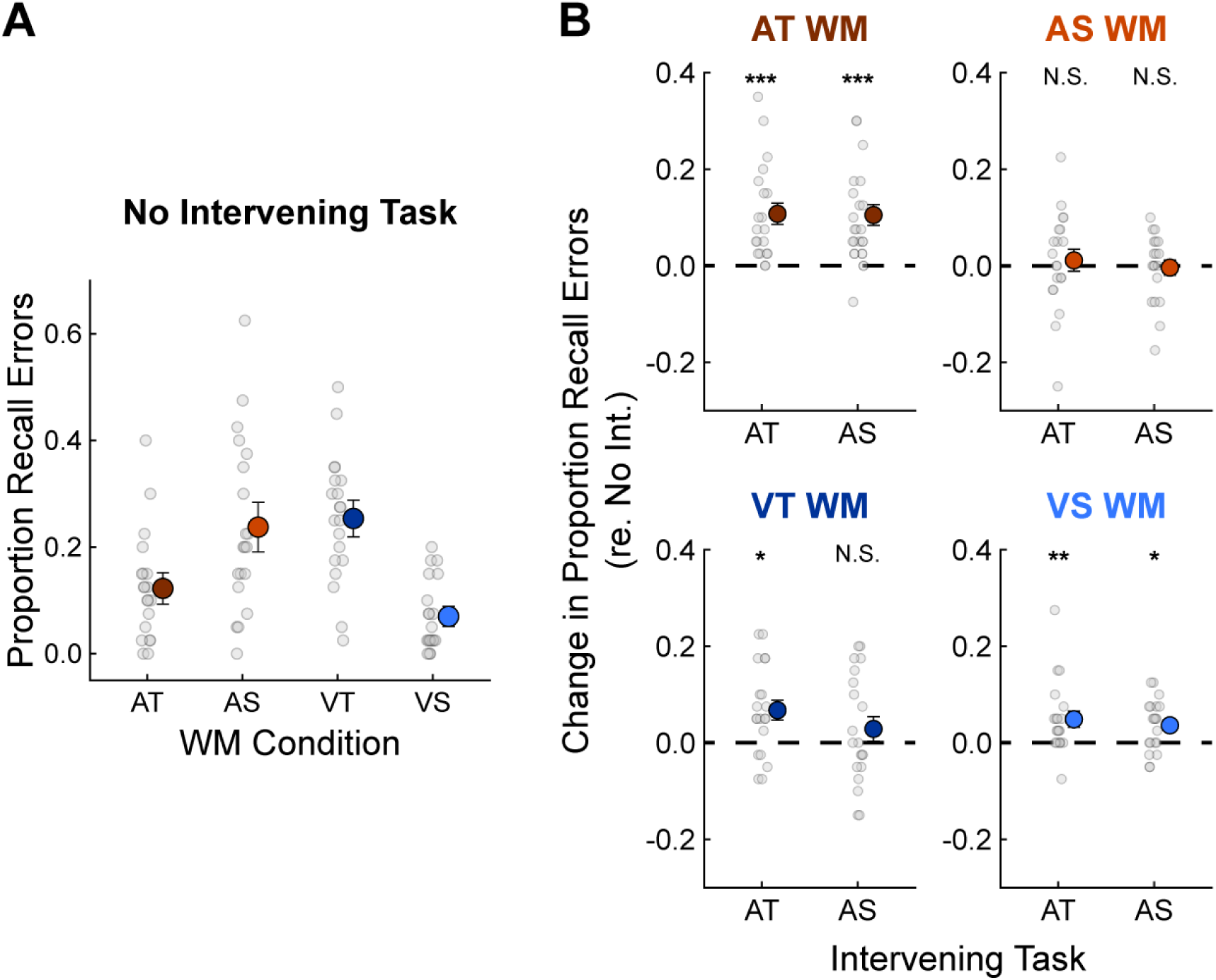
Working memory task performance. **A**, The proportion of trials on which the WM information was retrieved incorrectly is plotted for conditions without an Intervening task. Chance performance is at 0.5. **B**, The change in retrieval error rate is plotted for each WM and Intervening task combination relative to the no Intervening task conditions. The dashed line at 0 indicates no change in WM retrieval. Post-hoc comparisons were limited to differences between Intervening task conditions within each WM task. Asterisks indicate significant differences from the corresponding no Intervening task conditions (no significant differences were found between the two Intervening tasks). * = p < 0.05, ** = p < 0.01, *** = p < 0.001 after Bonferroni-Holm correction.

### Working Memory Task Performance

Participants were able to retrieve the information stored in WM at better-than-chance levels across all combinations of WM modality, WM domain, and Intervening task. These factors (including conditions with no Intervening task) served as fixed effects in a logistic mixed effects model of WM task retrieval errors. A three-way ANOVA conducted on the resulting model coefficients showed a significant interaction between WM task modality and domain (χ^2^(1,20) = 206.4, p < 10^−15^), reflecting overall lower error rates on the AT and VS WM tasks (Fig. 3A). In these conditions, the sensory modality was optimally suited for the information domain of the WM task. The ANOVA also revealed a significant three-way interaction between WM modality, WM domain, and Intervening task (χ^2^(2,20) = 19.0, *p =* 7.66·10^−5^), indicating different effects of the Intervening tasks depending on the WM condition. These effects were explored via post-hoc testing conducted separately within each WM condition, so that differences in WM retrieval could be attributable solely to interference from the Intervening tasks.

To visualize effects of the Intervening tasks on recall accuracy, Fig. 3B shows the *difference* between WM error rates in each Intervening task condition and the corresponding condition with no Intervening task. When the WM task was AT (top-left panel), both auditory Intervening tasks significantly impaired WM retrieval (p < 10^−6^ for both). When the WM task was AS, on the other hand, the Intervening tasks had no detectable impact on WM retrieval (top-right panel). We suspect that holding AT information in WM relied heavily on the auditory-biased WM network, leading to memory interference from the auditory Intervening tasks. In the AS WM task, on the other hand, sound locations to be remembered may have been, at least in part, mapped into a representation in the visual-spatial WM network, helping to protect this information from auditory interference (see Discussion).

Recall accuracy was relatively poor in the AS WM condition, which may have impaired our ability to detect effects of the Intervening tasks in this condition, as closer-to-chance performance left less room for performance to be impaired by the Intervening tasks (Fig. 3A). To explore this possibility, we built a linear model of the effects of the Intervening tasks (the difference in recall performance between conditions with and without an Intervening task) as a function of performance in the no Intervening task condition. There was no statistically significant relationship when the Intervening task was AS (p = 0.31); thus, the degree of interference from the Intervening task did not depend on baseline AS WM performance. For the AT Intervening task, the effect was marginally significant (F(1,18) = 4.71, *p =* 0.044), but this effect hinged on the one participant who performed at worse-than-chance levels in the No Intervening task condition; without this participant, the effect disappeared (F(1,17) = 0.54, *p =* 0.47). Thus, we conclude that the auditory Intervening tasks had minimal effect on participants’ ability to recall AS information from WM.

In some cases, visual WM retrieval was also modestly impaired by the auditory Intervening tasks. A significant increase in retrieval errors on the VT WM task was detected when the Intervening task was also temporal (*p =* 0.042; bottom-left panel of Fig. 3B). This might reflect domain-based WM interference with both tasks drawing on temporal processing resources, although no significant difference was found between the AT and AS Intervening task conditions. In the VS WM condition, both Intervening tasks increased retrieval errors (*p =* 0.003 for AT Intervening task, *p =* 0.044 for AS; bottom-right panel).

### Pupil Dilations

Both encoding information in WM and performing the auditory Intervening tasks led to reliable pupil dilations, shown in Fig. 4. In the absence of an Intervening task, pupil diameter increased throughout presentation of the stimuli to be encoded in WM (from −2 to 0 sec on the time axis), peaked about 700 ms after the offset of the final stimulus in the WM encoding phase, and then gradually declined throughout WM retention (Fig. 4A, left panel). Pupil size was significantly larger during the encoding phase of the AS WM condition compared to any of the other WM conditions.

**Figure 4:**
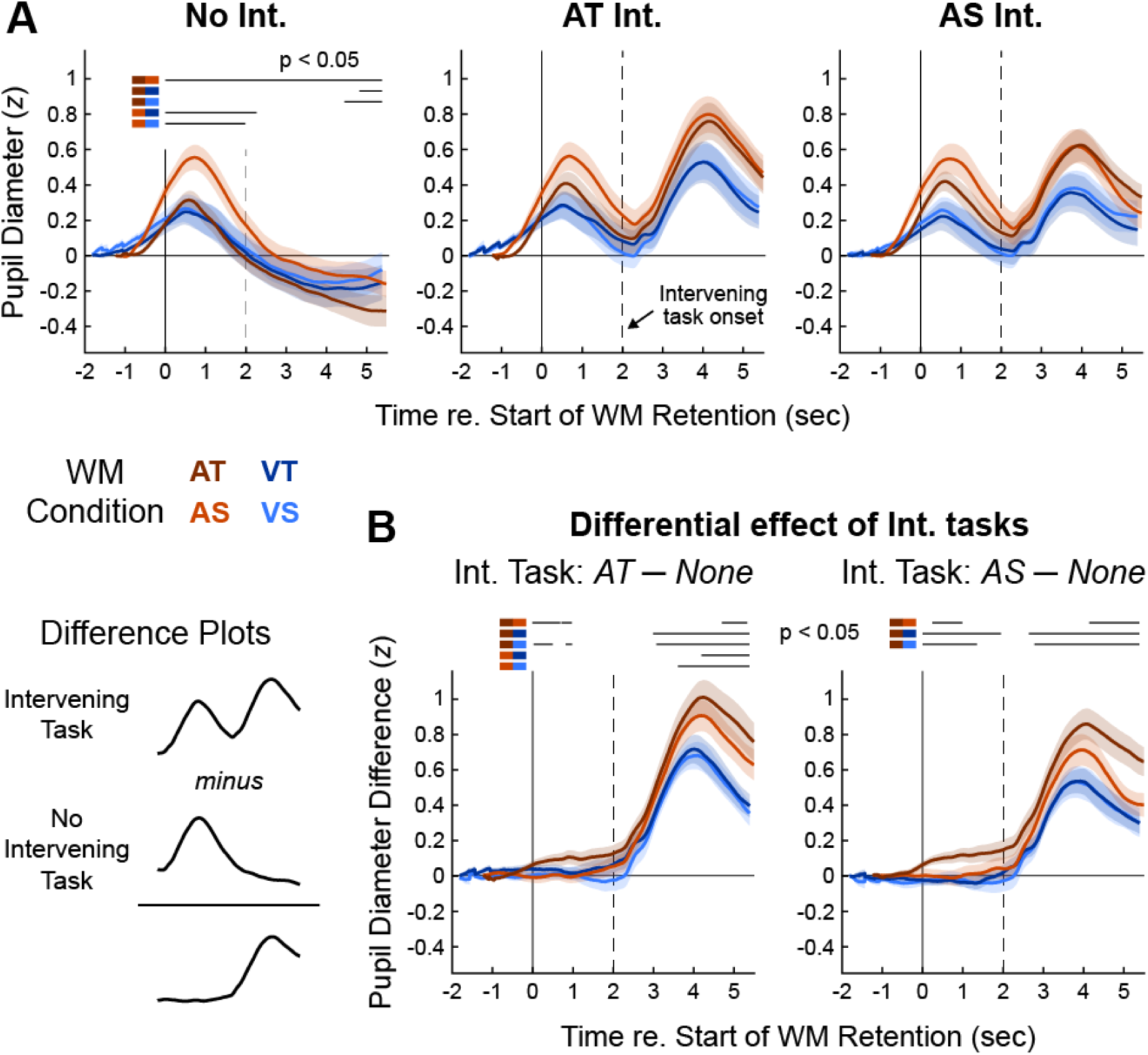
Pupillometry time courses. **A**, Grand average Z-scored pupil responses are shown in the No Intervening task condition (left) and elicited by physically identical stimuli in the AT (middle) and AS (right) Intervening tasks. Solid vertical lines represent the end of WM encoding and the start of WM retention, while dashed vertical lines indicate the onset of the Intervening task when present. Error clouds represent S.E.M. Horizontal black lines above the traces represent time regions of significant difference in permutation testing, with the two WM conditions being compared indicated by colors next to each significance line. **B**, The difference between pupil responses elicited in conditions with an Intervening task and responses in the corresponding WM conditions with no Intervening task.

There were also large secondary pupil dilations in conditions with an Intervening task, which started shortly after Intervening task onset but peaked around 4s into WM retention, approximately 1s after the Intervening task behavioral response. The amplitudes of these pupil responses were modulated by the information participants were holding in WM (Fig. 4A, middle and right panels). To better isolate the impact of the Intervening tasks on pupil diameter, we subtracted corresponding pupil traces with no Intervening task from each pupil trace with an Intervening task (Fig. 4B). Statistical differences between the pupil responses were assessed using non-parametric permutation testing, restricted to a time window spanning the WM retention phase (time zero and later in Fig. 4). We first averaged the differential pupil traces across the four WM conditions and tested for a difference between the two Intervening tasks (not shown). This confirmed that traces elicited by the AT Intervening task were larger overall than those elicited by the AS Intervening task (p < 0.05 for all time points between 3.7 sec after WM retention onset and the end of the WM retention window), matching the higher behavioral error rates on the AT Intervening task.

Next, we compared the differential pupil traces between the four WM conditions within each Intervening task (Fig. 4B), similar to our treatment of the Intervening task behavioral data. Differential pupil dilations elicited by both Intervening tasks were larger when the information held in WM was auditory than when it was visual, mirroring the modality-driven interference effects observed in the behavioral data. The largest differential pupil dilation occurred when the WM task was AT—significantly larger than when the WM task was AS—for both Intervening tasks. In the AT WM condition, the residual pupil traces were also elevated between 0 and 2 seconds from the start of WM retention (prior to the onset of the Intervening task), indicating a preparatory increase in effort in this WM condition. Combined, these results suggest that participants increased task effort both before and during the Intervening task when the information held in WM was AT, as both auditory Intervening tasks strongly interfered with recall in this WM condition. In the visual WM conditions, pupil dilations were insensitive to whether the WM task was temporal or spatial. This result is similar to the Intervening task error rates and indicates that interference was generally low when the two tasks were presented in different sensory modalities, even if they were matched in information domain.

We also analyzed average pupil size during the pre-trial baseline period, primarily to assess whether our baseline correction procedure may have biased the subsequent pupil dilation patterns. We conducted a separate two-way ANOVA for each Intervening task condition, with factors of WM modality and WM domain. In the AT Intervening task ANOVA, there was a significant main effect of WM modality (F(1,19) = 9.89, *p =* 0.005, partial-η^2^ = 0.045), with baseline pupil size tending to be smaller in the auditory than the visual WM conditions (Fig. 5). This is an intriguing result, as the AT Intervening task was relatively difficult and tended to interfere with auditory (particularly AT) WM. Smaller pupil size in the auditory WM conditions might indicate fatigue during these challenging blocks (Morad et al., 2000). However, there were no significant baseline pupil size effects in the AS Intervening task ANOVA (nor the no Intervening task ANOVA). Since the pattern of task-evoked pupil dilations was not qualitatively different between the AT and AS Intervening task conditions, it is unlikely that baseline correction biased the main results in any meaningful way.

**Figure 5:**
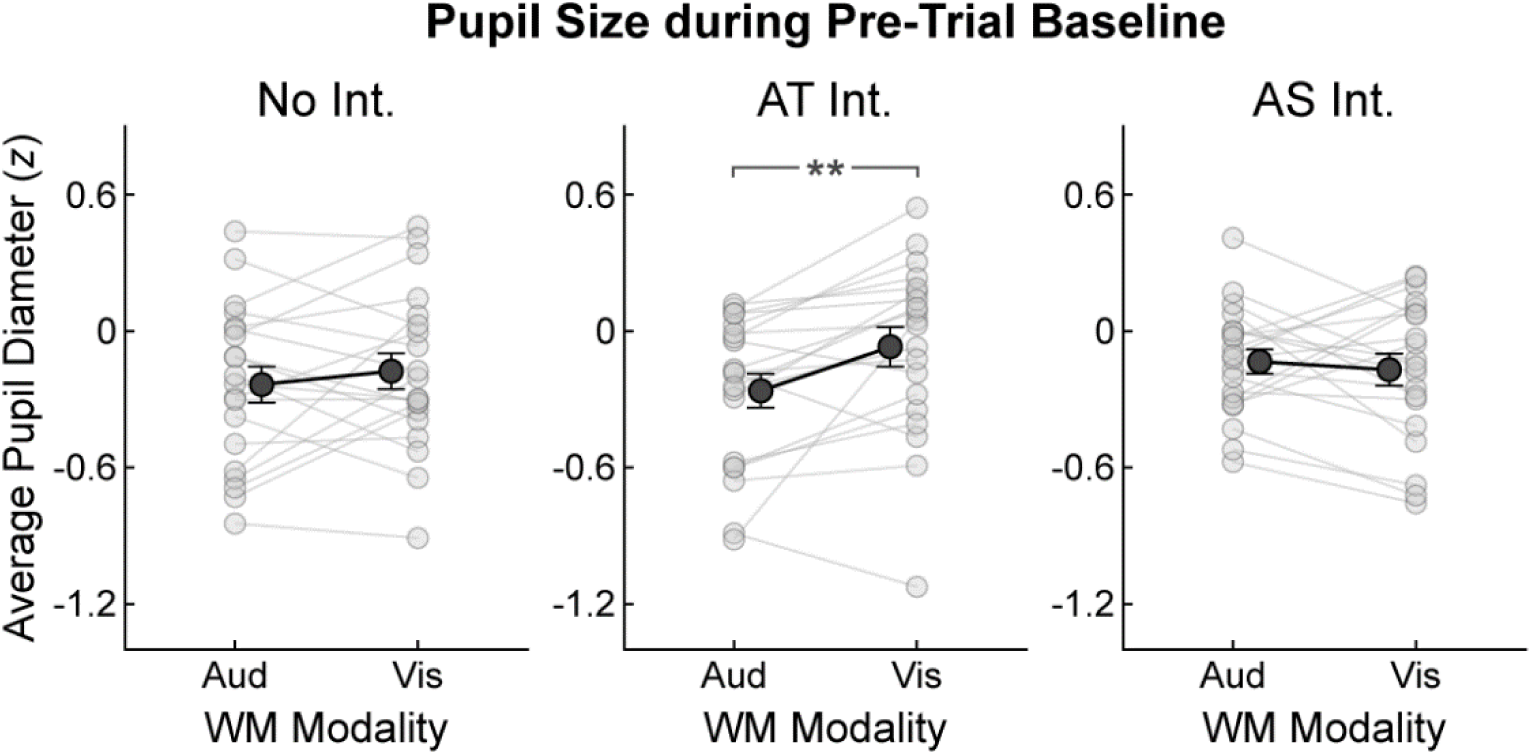
Baseline pupil size. Z-scored pupil size, averaged across the pre-trial baseline window, is shown for each Intervening task condition and WM task modality. Data are collapsed across WM domain, as this factor had no statistical influence on baseline pupil size. Light grey points and lines represent individual participants, dark grey points represent grand averages, and error bars represent S.E.M. ** = p < 0.01.

### Event-related potential (ERP) amplitudes

While the behavioral and pupillometry data revealed consistent effects of the modality and domain of the WM task, the domain of the Intervening task had relatively little impact on signatures of dual-task interference. The AT and AS Intervening tasks did require different types of auditory processing however, which could affect patterns of dual-task interference at the neural level. We next investigated this possibility in the EEG data.

We first examined event-related potentials (ERPs) elicited by each stimulus event in the WM and Intervening tasks (Fig. 6). ERPs elicited by stimuli in the encoding and probe sequences of the WM task were not systematically modulated by the Intervening tasks. However, the magnitude of the ERPs elicited by the Intervening task stimuli—particularly the first sound in the Intervening task sequence (highlighted in Fig. 6)—varied depending on the information participants were holding in WM. The Intervening task auditory ERPs had a somewhat atypical morphology, including a weak N1 and a strong component in the P2 time range (compare the Intervening task ERPs to auditory ERPs elicited by the WM task stimuli). For convenience, we will refer to this positive component as the “P2” and return to its interpretation in the Discussion.

**Figure 6:**
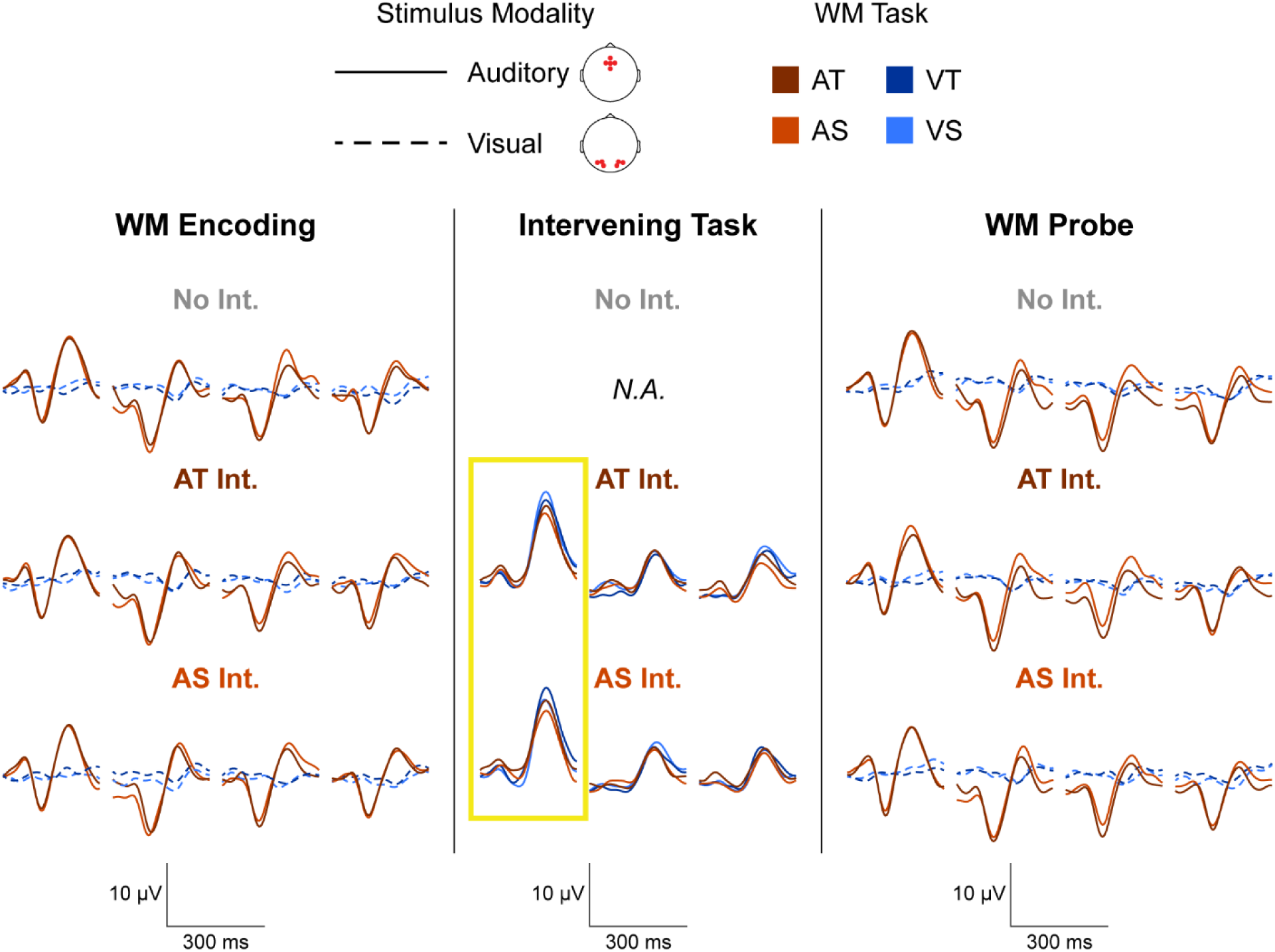
Event-related potentials. Grand-average ERPs are shown for each combination of WM task (colors), Intervening task (rows), and position in the stimulus sequence. The modality of stimulus presentation is shown by solid versus dashed lines. Intervening task onset ERPs, which were subsequently analyzed in greater detail, are highlighted by the yellow box. The top-left legend shows the channels averaged to produce ERPs for each stimulus modality: on the standard 10-20 layout, these were channels Fz, AFz, Cz, F1, and F2 for auditory stimuli and O1, O2, PO3, PO4, PO7, PO8 for visual stimuli. Axes below the data in each trial phase are shown for scale.

Both the modality and domain of the information stored in WM affected the amplitudes of the P2 component (Fig. 7, panels A and B). To quantify these effects, P2 amplitudes were calculated for each participant as the average of the ERP waveform between 170 and 230 ms post-stimulus across a cluster of fronto-central electrode sites (Fz, FCz, Cz, FC1, and FC2 on the standard 10-20 layout). These amplitudes were then submitted to a three-way ANOVA with explanatory factors of WM modality, WM domain, and Intervening task. This analysis revealed a significant main effect of WM modality (F(1,19) = 7.27, *p =* 0.014, η^2^ = 0.038) and a significant interaction between WM domain and Intervening task (F(1,19) = 10.83, *p =* 0.004, η^2^ = 0.005).

**Figure 7:**
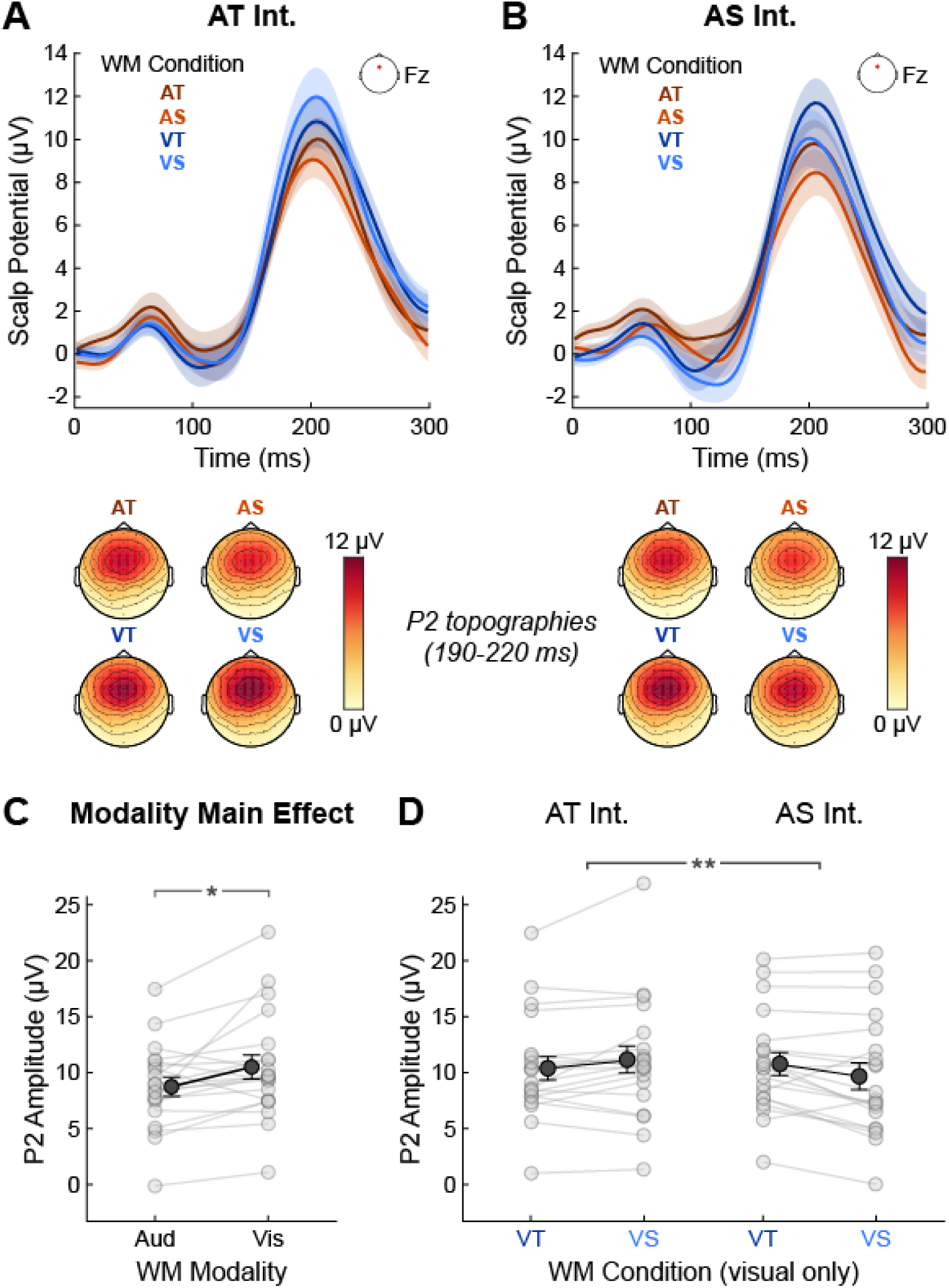
Intervening task onset ERPs. **A**, Grand average ERPs elicited by the first sound in the AT Intervening task at a fronto-central electrode site (Fz). Color represents the WM task, and error clouds represent S.E.M. Scalp topographies of the P2 responses are shown below the traces. **B**, Same as A, but for the AS Intervening task. The ERPs shown in A and B are those highlighted in Fig. 6. **C**, P2 peaks in the auditory and visual WM conditions, averaged across WM task domain and Intervening task. **D**, P2 peaks in the visual WM conditions to illustrate the interaction between WM domain and Intervening task.

The main effect of WM modality reflects the fact that P2 amplitudes elicited by both auditory Intervening tasks were larger when visual rather than auditory information was stored in WM (Fig. 7C). Although the three-way interaction involving WM task modality did not reach statistical significance, the significant interaction appears to have been driven predominantly by the visual WM conditions. We therefore split the P2 peak data into auditory and visual WM conditions, then conducted separate 2-way ANOVAs with factors of WM domain and Intervening task.

In the visual WM conditions, the interaction between WM domain and Intervening task was significant (F(1,19) = 10.77, η^2^ = 0.009, *p =* 0.004; beneath a Bonferroni-Holm adjusted alpha criterion to account for the two additional ANOVAs). Specifically, ERPs elicited by the AT Intervening task were slightly larger when visual-*spatial* information was stored in WM, whereas ERPs elicited by the AS Intervening task were larger when visual-*temporal* information was stored in WM (Fig. 7D). This provides evidence that P2 components of the ERPs were reduced when the two tasks required processing in the same information domain. However, in the auditory WM ANOVA, there was a significant effect of WM domain (F(1,19) = 5.21, *p =* 0.034, η^2^ = 0.015) but no interaction with Intervening task; although the significance of the WM effect did not survive Bonferroni-Holm adjustment of the alpha criterion, the trend was for ERPs elicited by either Intervening task to be larger when AT information was stored in WM than when the WM task was AS.

### Alpha-band oscillatory activity

Changes in oscillatory activity in the alpha band (8-13 Hz) have been linked to maintaining information in WM (Bastiaansen et al., 2002; Obleser et al., 2012). We investigated whether this neural signature of WM maintenance was affected by the presence of an Intervening task, as well as the degree of similarity in sensory modality and information domain between the Intervening and WM tasks. Figure 8A shows the power spectrum during the pre-trial baseline period at a posterior electrode site (Pz), with a strong peak in the alpha range, validating that this oscillatory signature occurred in the present data. An anterior electrode site is also shown, with expected oscillatory power in the 4-7 Hz range (frontal theta oscillations).

**Figure 8:**
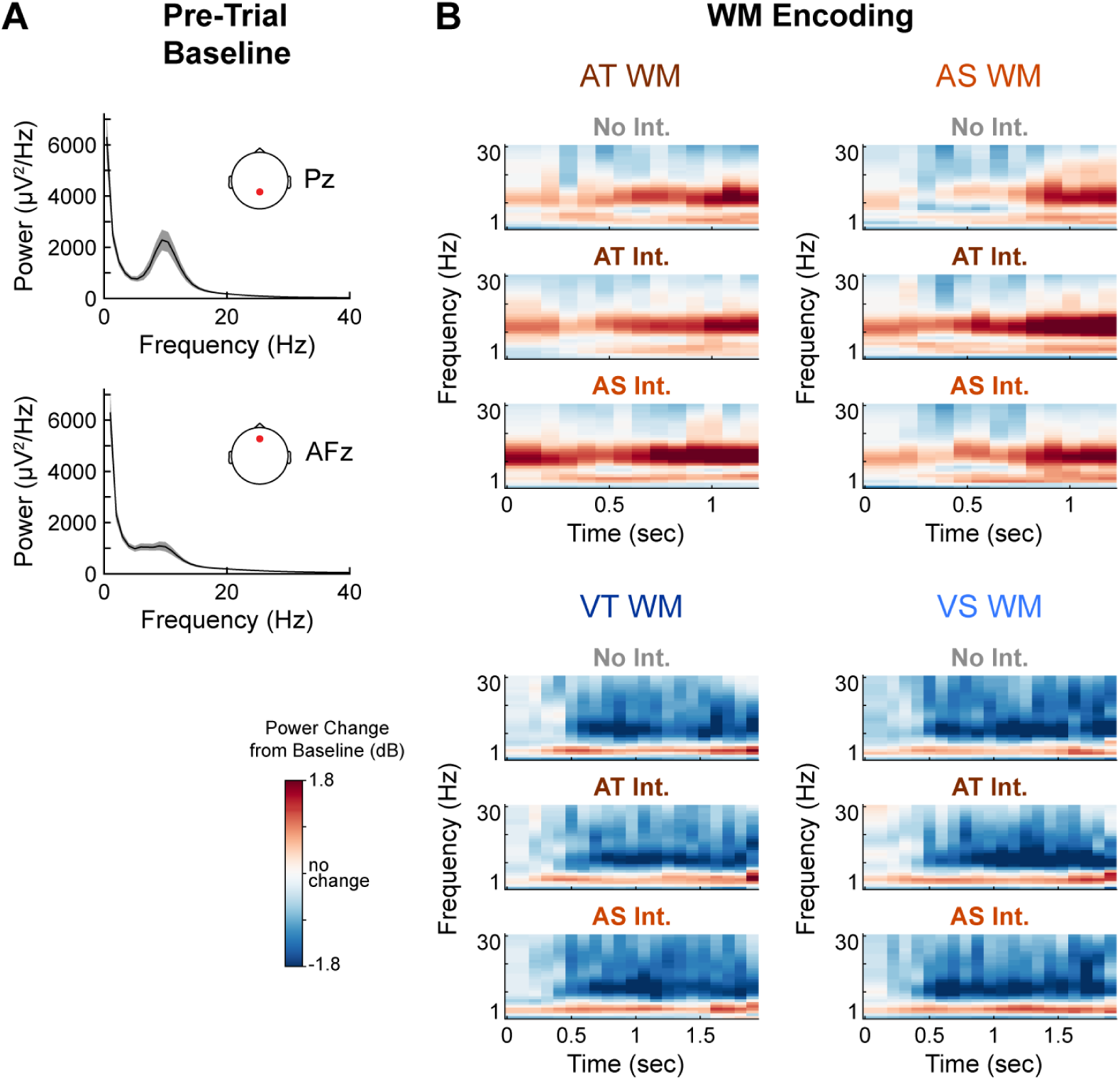
Oscillatory power in the baseline and during WM encoding. **A**, Grand average power spectra during the pre-trial baseline, plotted at a posterior and an anterior electrode site. Error clouds represent S.E.M. **B**, Grand average oscillatory power during WM encoding in the AT, AS, VT, and VS WM conditions. Data are shown as change relative to the pre-trial baseline, in dB. Because the length of the encoding window differed based on inter-stimulus intervals, responses of different lengths were aligned to the end of the encoding phase prior to averaging. Similarly, because the long interval was longer for visual WM than auditory WM, the time bases differ between the top and bottom panels.

During WM encoding, alpha power steadily increased relative to baseline in the auditory WM conditions and decreased relative to baseline in the visual WM conditions (Fig. 8B). These gross patterns of alpha activity persisted into the WM retention window. Preliminary analysis of alpha power across the retention window using cluster-based permutation testing revealed no significant effects of WM task domain, so data were collapsed across this factor and subsequently analyzed based on WM task modality and the Intervening task.

In the auditory WM tasks, alpha power remained elevated relative to baseline throughout the memory retention window when no Intervening task was present (Fig. 9A, top panel). The onset of both the AT and AS Intervening tasks suppressed these ongoing alpha oscillations (middle and bottom panels). We examined this effect by conducting cluster-based permutation tests on the alpha power time courses in two time regions of interest: one spanning the Intervening task itself (2 to 3.5 seconds into WM retention, including the Intervening task behavioral response), and another from 3.5 to 5 sec, during which participants needed to start retrieving the information stored in WM in anticipation of the WM task probe sequence. During both of the Intervening tasks (the earlier window), alpha power was significantly reduced relative to the no Intervening task condition (AT Intervening task, *p =* 0.005; AS Intervening task, *p =* 0.003; Fig 9B). With the statistical threshold for generating the initial clusters set at *p =* 0.01, no differences were found between alpha power in the two Intervening tasks. However, relaxing this threshold to *p =* 0.05 (not shown) revealed an additional late cluster showing an alpha power difference between the AT and AS Intervening tasks (significant between 3.14 and 3.45 sec; *p =* 0.026). This suggests that the alpha power reduction lasted longer when the Intervening task was AT than when it was AS, probably because participants could sometimes make their perceptual judgment earlier in the AS Intervening task by comparing the first sound location to the second (without using the third). No significant clusters were found in the later (3.5-5 sec) time window in the auditory WM conditions.

**Figure 9:**
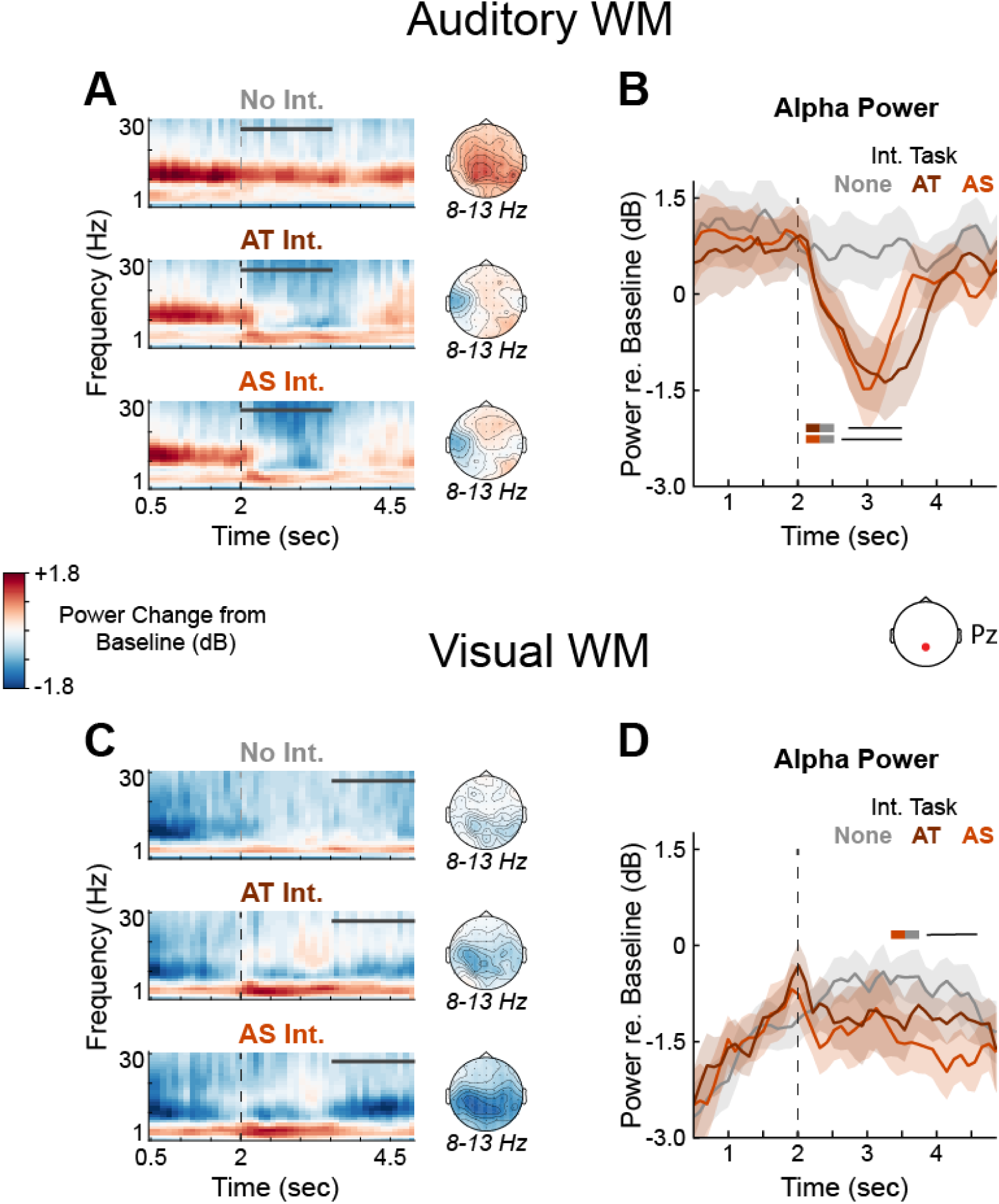
Alpha power during WM retention. Grand-average time-frequency responses are shown at a parietal electrode site (Pz) during the memory retention window for each of the three Intervening tasks in auditory (**A**) and visual (**C**) WM conditions. Responses are averaged across WM domain and shown as dB change relative to the average pre-trial baseline period across conditions. Vertical dashed lines represent Intervening task onset when present, and horizontal grey bars represent the time regions in which significant effects were found in cluster-based permutation testing. The first and last 500 ms of the retention window were excluded to limit power contributions from responses evoked by the WM task stimuli. B, Grand average alpha power time courses are also shown for the auditory (**B**) and visual **(D)** WM conditions. Power time courses for each participant were calculated at their peak alpha frequency ±1 Hz. Error clouds represent S.E.M. Black horizontal bars indicate the temporal extent of significant clusters in cluster-based permutation testing for the comparisons indicated by colored boxes to the left.

In the visual WM conditions without an Intervening task, alpha power gradually increased from the suppressed levels during WM encoding to near-baseline levels during WM retention (Fig. 9C, top panel). On trials with an Intervening task, alpha power become suppressed again after the conclusion of the Intervening task (bottom panels). We examined the alpha power time courses using cluster-based permutation testing restricted to the same two time windows as in the auditory WM conditions. Unlike the auditory conditions, no significant differences were observed during the Intervening tasks. However, in the later window, when participants needed to transition from the Intervening task to WM recall, alpha power was significantly reduced when the Intervening task was AS relative to when there was no Intervening task (*p =* 0.011; Fig. 9D). Alpha power did not differ either between the AT and no Intervening task conditions or between the AT and AS Intervening tasks.

Finally, we examined frontal theta band (4-7 Hz) oscillatory activity, as increased theta power has been implicated in paradigms that require task-switching (Cunillera et al., 2012; López et al., 2019; Sauseng et al., 2006). The presence of either Intervening task caused an increase in frontal theta power throughout the memory retention window (not shown). In all but one of the conditions with an Intervening task, cluster-based permutation tests revealed at least one time-channel cluster in which theta power was significantly elevated relative to the corresponding no Intervening task conditions (p < 0.05 for all clusters). In the remaining condition (AT WM, AS Intervening task), a similar increase in theta power trended toward significance (*p =* 0.053). Although ERPs can manifest as activity in this frequency range as well, the increase in theta power was observed both preceding and well after the Intervening task, meaning it cannot easily be explained by evoked responses (grand average, condition-specific ERPs were also subtracted from the time series prior to time-frequency analyses). The broad presence of increased theta power across conditions with an Intervening task suggests that switching from WM encoding to perceptual processing drove this neural signature, but it generally did not differ based on the condition of either task.

## Discussion

In this study, we investigated interference between perceptual processing and WM maintenance while varying similarity in sensory modality and information domain. Convergent evidence from behavior and pupillometry (indexing cognitive effort) showed that the highest degree of interference occurred when the two tasks drew upon shared cognitive resources. Specifically, when both tasks were auditory-temporal (AT), error rates on the Intervening task were highest, WM retrieval was poorest, and pupil dilations were largest. In contrast, when the auditory Intervening tasks were paired with a visual (visual-temporal, VT or visual-spatial, VS) WM condition, behavioral interference was relatively low and pupil size was smallest. In these conditions with reduced competition for shared neural resources, subtler patterns of interference related to the task domain were nonetheless present in the EEG data. Results from behavior and pupillometry indicated an intermediate level of interference from the auditory Intervening tasks when the WM task was auditory-spatial (AS); this condition will receive special attention below, as we expected maintaining AS information in WM to draw on both auditory- and visual-biased control networks, leading to more nuanced patterns of interference.

### Interference patterns support modality and information domain specializations of lateral frontal control networks

The design of this experiment was inspired by recently characterized attention and WM networks involving human lateral frontal cortex (LFC). Subregions of the LFC preferentially contribute to auditory or visual processing, but are also recruited by stimuli in the non-preferred modality depending on whether the task is temporal or spatial. In this study, our goal was to examine the influence of sensory modality and information domain on dual-task interference, informed by the response profiles of these networks. We did not use neuroimaging methods (such as fMRI) that would allow us to directly relate dual-task interference to activity in subregions of the LFC, but our WM task conditions were derived from tasks shown to recruit these networks in fMRI experiments (Michalka et al., 2015). Thus, we are confident that the AT WM condition strongly engaged the auditory-biased network, the VS WM condition strongly engaged the visual-biased network, and the AS and VT WM conditions recruited both networks. However, recruitment of these LFC networks is also asymmetric; auditory processing activates the visual-biased network to a greater extent than visual processing activates the auditory-biased network (Noyce et al., 2017, 2022). The AS WM condition and AS Intervening task are therefore of particular interest in the present study, as we expected that auditory information in these tasks might ultimately be mapped to, and maintained in, the visual-biased network. One prediction from this is that perceptual processing that taxes the auditory-biased network should be less affected by holding AS information than holding AT information in WM, as the AS information can be represented (at least partially) in the complementary visual-spatial network. In line with this prediction, participants made fewer errors on the AT Intervening task (which we expected to load exclusively onto the auditory-biased network) when the WM task was AS than when it was AT. The AS Intervening task, on the other hand, may have recruited both networks to some extent, leading to similar levels of behavioral interference from a WM condition that overloaded the auditory-biased network (AT WM) and one that increased the overall distributed load across the two networks (AS WM).

We initially expected that interference effects between the two tasks would be symmetrical—that is, that effects of the Intervening task on WM recall would be similar to effects of the WM task on Intervening task performance. However, whereas the WM conditions differentially affected Intervening task performance, the two Intervening tasks interfered with WM recall roughly equally. This likely relates to important differences in processing demands between the AS WM task and the AS Intervening task. To perform the AS WM task, participants needed to remember absolute auditory stimulus locations mapped onto physical space. This may have placed more demand on the visual-biased LFC network than the AS Intervening task, which required only an immediate, relative judgment about auditory spatial positions within each trial, and thus did not need to be stored in memory. While the AS Intervening task certainly involved a form of spatial processing that could interfere with other spatial tasks, it seems likely that it primarily taxed the auditory-biased LFC network, similar to the AT Intervening task. This would explain why both Intervening tasks strongly interfered with AT, but not AS, WM recall, and why pupil dilations elicited by both Intervening tasks showed the highest effort with AT information in WM. An important (if unintended) point to draw from this is that two tasks with ostensibly similar processing demands could in actuality draw upon different neural resources; in this case, not all spatial tasks are created equal.

Another noteworthy aspect of the results presented here is that the processing costs of loading onto a single network were greater than the costs of switching between networks. This outcome is not entirely obvious, as task-switching also incurs a processing cost (Arnell & Jolicœur, 1999; Chun & Potter, 2001; Hsieh & Allport, 1994; Meiran et al., 2000). Our auditory Intervening tasks did modestly impair visual WM retrieval, consistent with a behavioral cost of switching between the complementary networks. However, this effect was relatively weak compared to the behavioral (and physiological) costs when the tasks loaded onto shared network resources. Thus, the ability to simultaneously leverage complementary attention and WM networks appears to confer a processing benefit, which may be especially relevant in settings with simultaneous inputs from multiple sensory modalities.

In this study, we only used auditory Intervening tasks to limit the influence of visual stimuli on the pupil response. The lack of visual Intervening tasks resulted in an upper limit on how much load could be placed on the visual-biased control network, as there was no condition involving competing visual tasks. Future studies (even if they are only behavioral) should implement visual Intervening tasks in a paradigm like this one to examine whether dual-task conditions that load exclusively on the visual-biased network (i.e., two visual-spatial tasks) also result in elevated interference. Domain-based patterns of interference might be more muted in such an experiment, however, as recruitment of the auditory-biased LFC network by visual-temporal tasks is relatively modest (Noyce et al., 2017).

### Unique contributions from pupillometry

The pupillometry data in this study revealed details about the magnitude and time course of interference that could not be discerned from behavioral responses alone. For instance, the AS Intervening task was easy enough that a majority of participants did not make errors on it in any WM condition. However, substantial differences in the amplitude of pupil dilations elicited by this task suggest that AS perceptual processing and WM maintenance did cause mutual interference, which participants overcame to achieve near-ceiling performance. Indeed, a major benefit of pupillometry is that it can reflect differences in the effort required to reach a certain performance level in the absence of behavioral differences (e.g., McGarrigle et al., 2017; Ohlenforst et al., 2017; Winn & Teece, 2021). Pupil responses have also provided useful insights into the temporal dynamics of effort allocation in cognitive processes that unfold through time, such as decision making (Satterthwaite et al., 2007) and handling talker variability (Lim et al., 2021). In this study, when the WM task was AT, pupil size increased prior to the onset of the Intervening task stimuli. This suggests a preemptive modulation of effort, perhaps reflecting the need to preserve the WM trace in anticipation of auditory interference.

The main pupil responses of interest in this study were those elicited by Intervening tasks under different types of WM load. These pupil dilations started shortly after Intervening task onset, but pupil size peaked after the Intervening task behavioral response and remained elevated throughout WM retention. Thus, these responses likely represent the combined effects of performing the Intervening tasks under WM load and recalling information from WM after auditory interference. This could explain why they are in accord with some aspects of both the WM task and Intervening task behavioral data. Consistent with WM recall, pupil responses were largest in the AT WM condition, and the pattern across WM conditions was similar between the two Intervening tasks. Consistent with Intervening task performance, pupil dilations were intermediate in the AS WM condition (when analyzed as the difference from the no Intervening task conditions), hinting that recruitment of the visual-biased network to store the AS WM information reduced the effort required to overcome auditory interference.

Many studies have shown that pupil size scales with task difficulty (Bijleveld et al., 2009; Kahneman & Beatty, 1966; Porter et al., 2007; van der Meer et al., 2010; Zhao et al., 2019), at least up to the point that the task becomes so difficult that participants disengage from it (Granholm et al., 1996). In line with this, overall pupil size (across WM conditions) was larger in response to the more difficult AT Intervening task than the AS Intervening task. However, most of the patterns of pupil size we observed cannot be explained by task difficulty alone. For instance, pupil responses after the Intervening tasks were largest when participants had AT information in WM, despite more accurate recall in this condition than in the AS or VT WM conditions. A parsimonious explanation, consistent with the behavioral data, is that these pupil responses were more tightly linked to patterns of modality- and domain-based interference than the combined difficulty of the two tasks.

Interestingly, the initial pupil response during WM encoding was substantially larger in the AS WM condition than any of the other conditions. Participants did make relatively frequent behavioral errors in the AS WM condition, suggesting a task difficulty effect, but a similar performance difference between VT and VS recall (recall errors were more frequent in the VT condition) was not reflected in the pupil data. Since the pupil data did not simply mirror task performance, the larger pupil dilations in the AS WM condition suggest that encoding AS information in WM is especially effortful, perhaps due to the simultaneous recruitment of both complementary control networks. In other contexts, types of processing that require extra effort ultimately yield perceptual benefits. For instance, using context to fill in masked words supports speech understanding but requires effort (Winn & Moore, 2018), and integrating visual information in speech processing improves intelligibility (Sumby & Pollack, 1954) but may be effortful (Strand et al., 2020). Similarly, encoding AS information in WM appears to require additional effort, but leveraging a visual-spatial framework to encode this information likely results in the most robust memory storage.

### Insights from the neural data

The fact that the Intervening tasks were always auditory limited the overall load on the visual-biased control network in this study. In the visual WM conditions, auditory interference did not rise to the level of affecting behavioral performance or effort requirements. In accord with the strongest interference occurring when both tasks were auditory, the auditory Intervening tasks elicited larger ERPs with visual as compared to auditory information held in WM. We posit that this indicates greater resource availability for auditory perceptual processing when participants were under a visual rather than auditory WM load. However, an alternative explanation could be that stimuli presented during auditory WM encoding caused adaptation of neural responses, resulting in suppressed amplitudes of auditory ERPs elicited by the onset of the Intervening task. We do not suspect this was the case for two reasons. First, although the two-second gap between the end of WM encoding and Intervening task onset was close enough that adaptation could theoretically have had some effect on the ERPs (Budd et al., 1998), neural responses would have likely been reset by the acoustic differences between the WM and Intervening task stimuli, as observed in auditory oddball paradigms (Paavilainen, 2013). Second, adaptation cannot account for differences in ERP amplitudes driven by differences in information domain (discussed below), as the stimuli were physically identical across task domain conditions.

Intervening task ERPs were largest when the two tasks differed in both sensory modality *and* information domain, even though similarity in information domain did not affect behavioral performance or pupil size in the visual WM conditions. Importantly, the VT and VS WM tasks used physically identical visual stimuli, and so differences in ERP amplitudes between these WM conditions cannot be attributed to bottom-up stimulus salience. Instead, these results indicate that the most neural resources were available for Intervening task processing when the two tasks differed in both modality and domain, leading to the largest evoked responses. This provides neural evidence that the WM control networks engaged by this task are organized not only according to the sensory input channel, but also the domain of the information being represented.

Within the auditory WM conditions, no such pattern of domain-based interference was found. Instead, ERPs elicited by both Intervening tasks tended to be slightly larger with AT than AS information held in WM. One possible explanation relates to the particularly high degree of effort required to encode AS information in WM, discussed in the previous section. If AS WM encoding consumed a disproportionate amount of domain-general cognitive resources also needed for attention in the Intervening task, ERP amplitudes in this WM condition may have been generally suppressed.

The primary ERP component that was modulated by our task manipulations peaked around 200ms post-stimulus, and we have therefore referred to it as the “P2”. However, since the Intervening task stimuli were preceded by either visual or acoustically distinct auditory stimuli, another possibility is that the large ERP positivity was actually the early phase of the novelty-P3 response, elicited by novel or unexpected sensory stimuli (Escera & Corral, 2007; Friedman et al., 2001; Squires et al., 1975). The novelty-P3 can have a peak latency as early as 220ms and a scalp distribution broadly consistent with the responses observed here (Escera et al., 1998). In this case, ERP differences in the visual WM conditions could imply a novelty response to a change in the information domain of the task. However, ERPs elicited by the second and third Intervening task stimuli also had this weak N1/strong P2 morphology (see Supplemental Fig. 2), less consistent with the novelty-P3. Alternatively, differences in auditory ERP morphology between the WM and Intervening tasks may have resulted from acoustic differences between the stimuli (tone complexes in the WM task, white noise bursts in the Intervening task). The data in Matsuo et al. (2019) showed that P3 amplitudes were larger in ERPs elicited by white noise bursts than pure tones, and parallels can be drawn between Intervening task ERPs in this study and the cortical responses to noise-like fricative phonemes in Khalighinejad et al. (2017).

Alpha oscillatory power was modulated by the type of information being held in WM and interrupted by the Intervening tasks. The largest alpha power changes from baseline were found at central-parietal electrode sites. Alpha activity with this scalp distribution has been linked to maintaining various types of information in WM, including visually presented letters and numbers (Jensen et al., 2002; Klimesch et al., 1999; Schack & Klimesch, 2002), shapes (Herrmann et al., 2004; Johnson et al., 2011), spatial information (Bastiaansen et al., 2002), and auditory stimuli (Lim et al., 2015; Obleser et al., 2012; Vogt et al., 1998). In the auditory WM conditions, alpha power was elevated during WM encoding and maintenance, but ongoing oscillations were interrupted by the auditory Intervening tasks. This is consistent with previous work showing that irrelevant visual stimuli interrupt alpha power during retention of both auditory and visual information in WM (Hakim et al., 2020; Mishra et al., 2013). In the visual conditions, alpha power was suppressed relative to baseline during WM encoding, likely because participants needed to attend task-relevant visual stimuli (Foxe & Snyder, 2011; Jensen & Mazaheri, 2010). This alpha suppression returned after participants performed the AS Intervening task, perhaps due to spatial interference in the visual-biased network. The spatial interference from the AS Intervening task suggested by the alpha and ERP data may be momentary in nature, as the AS Intervening task did not require mapping stimulus positions into memory. Such ephemeral interference required the temporal sensitivity of EEG to detect in this paradigm, but it may have meaningful processing consequences in real-world environments with many simultaneous sensory inputs and competing attentional and goal-oriented demands.

## Conclusions

In a dual-task paradigm designed to rely on auditory- and visual-biased attention and WM networks, we found behavioral, autonomic, and electrophysiological signatures of interference when the two tasks drew upon shared neural control resources. Specifically, when perceptual and working memory tasks were matched in both sensory modality and information domain, behavioral performance was worst, pupil dilations were largest, and ERP amplitudes were suppressed—all indicating strong dual-task interference. These results support “resource-specific” theories of WM maintenance and indicate that WM control networks are segregated not only on the basis of the sensory information channel, but also based on whether the task requires spatial or temporal information processing. The bidirectionality of interference effects between the two tasks suggests that WM maintenance and perceptual processing relied on shared neural resources in this study, most likely including sensory-biased control networks in the lateral frontal cortex. Future neuroimaging studies should expand this paradigm to explore the dynamics of how LFC control networks support the interplay between WM maintenance and ongoing perceptual processing.

## Acknowledgements

We would like to thank Jessica Tin and Yaminah Carter for their assistance with data collection and Matt Winn for discussions about interpreting the temporal dynamics of pupillometry data. The authors declare no competing interests. This work was supported by the Office of Naval Research (Grant N000141812069) and the National Institute on Deafness and Other Communication Disorders (Grant T32-DC000038).

